# Molecular determinants for Rous sarcoma virus intasome assemblies

**DOI:** 10.1101/2023.03.02.528060

**Authors:** Sibes Bera, Ke Shi, Hideki Aihara, Duane P. Grandgenett, Krishan K. Pandey

## Abstract

Integration of retroviral DNA into the host genome involves formation of integrase (IN)-DNA complexes termed intasomes. Here, we report the single-particle cryo-EM structure of the Rous sarcoma virus (RSV) strand transfer complex (STC) intasome produced with IN and a preassembled viral/target DNA substrate. The STC structure had an overall resolution of 3.36 Å and 3 Å in the conserved intasome core (CIC) region. Our structure demonstrated the flexibility of the distal IN subunits relative to the IN subunits in the CIC, similar to previously shown with the RSV octameric cleaved synaptic complex (CSC) intasome produced with IN and viral DNA only. An extensive analysis of higher-resolution STC structure helped in identification of nucleoprotein interactions important for intasome assembly. Using structure-function studies, we determined the mechanisms of several IN-DNA interactions critical for assembly of both RSV intasomes. We determined the role of IN residues R244, Y246 and S124 in CSC and STC intasome assemblies and their catalytic activities, demonstrating differential effects. Taken together, these studies advance our understanding of different RSV intasome structures and molecular determinants involved in their assembly.

## Introduction

Integration of retroviral DNA into the host genome, an essential step in retroviral replication, is mediated by the viral integrase (IN). Upon retrovirus infection, the RNA genome is converted to double-stranded DNA by viral reverse-transcriptase (RT). Both the IN and RT are closely associated with the viral genome within the virion core particle. IN binds to both ends of the linear viral DNA and removes two nucleotides from the 3’-ends adjacent to conserved CA dinucleotides exposing the reactive 3’-OH group. In the integration reaction, termed strand transfer, IN joins the processed viral 3’-OH ends with a scissile phosphate on the target DNA strands though a S_N_2-type nucleophilic substitution. This joining to the host DNA occurs in a staggered manner separated by 4 to 6 bp, a unique characteristic of different retroviral species (1).

IN from Rous sarcoma virus (RSV), HIV-1, and related retroviruses share a three-domain organization, namely a catalytic core domain (CCD) flanked by amino (NTD) and carboxy-terminal domains (CTD) (2). The NTD folds into a three-helix bundle and contains the conserved HHCC motif which coordinates a Zn^2+^ ion. The CCD is the most conserved domain among retroviruses and contains the active site comprising the DDE (Asp-Asp-Glu) motif which is responsible for catalytic activity. The CTD adopts an SH3 fold and is the least conserved domain. This domain may be the primary factor for producing different oligomeric forms of nucleoprotein complexes observed with different retroviral INs. The C-terminal tail region which extends beyond the CTD is flexible and has not been resolved in any IN structure. The tail region (17 aa) in RSV IN appears to have a major role in forming specific IN-DNA complexes termed intasomes. Truncations of the C-terminal tail favors the accumulation of a precursor tetrameric intasome en route to the mature octameric CSC intasome (3–5).

Purified IN exists in different oligomeric forms ranging from monomers to dimers to tetramers and higher order species. Virion-derived or recombinant RSV IN is predominantly dimeric (6) while recombinant HIV-1 IN is either monomeric, tetrameric or a mixture thereof (7–11). The oligomeric forms of free IN or IN associated with viral RNA or DNA in vivo remains unknown. IN multimerizes onto viral DNA ends to produce a series of complexes in the concerted integration pathway in vitro (12). IN coupling two 3’-OH processed viral DNA ends is referred as the cleaved synaptic complex (CSC) which upon binding to and integration into the target DNA is referred as the strand transfer complex (STC). Collectively, the CSC and STC and their intermediate complexes are termed intasomes. In general, most intasome assemblies observed in vitro suggest DNA-mediated tetramerization of the predominant IN species observed in solution. Monomeric PFV IN assembles tetrameric intasomes (13) while dimeric α-and β-retroviral INs form octameric intasomes (14, 15). Lentiviral INs like HIV-1, MVV, and SIV are mostly tetrameric and have been observed or expected to produce hexadecameric intasomes (16–19).

Intasome assembly mechanisms are not clearly defined. We used RSV IN as a model system to determine its intasome structures and to understand their assemblies and associated functions. We hypothesize there are key intermediates in the intasome assembly pathway with IN multimers and viral DNA that subsequently binds target DNA resulting in integration. Our previous studies demonstrated that an RSV tetrameric CSC intasome is the precursor to mature octameric CSC intasome (4, 5). The ability of RSV IN to assemble the catalytically active octameric intasome from its tetrameric precursor is unique among studied retroviral systems.

Previously, we determined the structure of RSV octameric CSC intasome by cryo-EM (20). The 4 proximal IN subunits complexed to the viral DNA form the conserved intasome core (CIC) (12) while the 4 distal subunits showed dynamic flexibility. We hypothesized that this flexibility could be stabilized by target DNA binding in the STC intasome, as shown in its X-ray crystal structure at 3.86 Å resolution (15). Cryo-EM captures DNA-protein complexes in their more native dynamic form while X-ray crystallography captures these complexes in a single more rigid conformation assisted by the crystal lattice. To resolve any potential structural bias, we determined the structure of RSV STC intasome in its native state by cryo-EM at an improved overall resolution of 3.36 Å. We carried out site-directed mutagenesis of selected IN residues to determine their effect on different intasome assemblies and catalytic functions. Collectively, these data advance the structural understanding of different RSV intasomes and molecular determinants involved in their assemblies.

## Results

### Cryo-EM structure of the RSV STC

STC intasomes were assembled using a DNA substrate that contains viral DNA covalently joined to target DNA (Fig. 1A). Upon integration, RSV produces a 6 bp host-site duplications at both ends of the cellular integration site. We used a similar 6 bp cellular sequence covalently linked at the 3’-end to the conserved CA residue at the viral LTR ends, thus mimicking the integration product. STC intasomes were purified by size-exclusion chromatography (SEC) using Superdex 200 Increase column (10 x 300 mm) (Fig. 1B). Unlike most other retroviral intasomes, we did not observe higher order species of STCs. RSV IN produced predominantly homogeneous octameric STCs (Fig. 1B). Similarly, homogeneous mature octameric CSC intasome is assembled with RSV IN and viral DNA (20). Top fractions of STCs from the SEC column were used for vitrification and cryo-EM imaging on Titan Krios microscope using Falcon 4 camera (*Table S1*). We determined the structure of RSV STC by single particle cryo-EM (Fig. 1C, *Fig. S1*, *Table S1*). We used cryoSPARC (21) to perform iterative 2D and 3D classification on our dataset of 3297 movies. We subsequently obtained a 3D map at 3.36 Å resolution from a stack of 141,428 particles (Fig. 1C). The CIC region that contained the viral and target DNA interactions within IN active sites had a resolution of ∼ 3 Å (*Fig. S1*). The cryo-EM structure demonstrated the presence of four IN dimers bound to the two STC DNA substates, modeled in Fig. 1D. The proximal subunits which have extensive interactions with DNA and the CTD region of distal subunits both had clear density and were well resolved. However, the distal NTD-CCD regions were resolved at a lower resolution probably due to dynamic flexibility observed also in this region with RSV CSC intasomes (20).

**Fig. 1.**
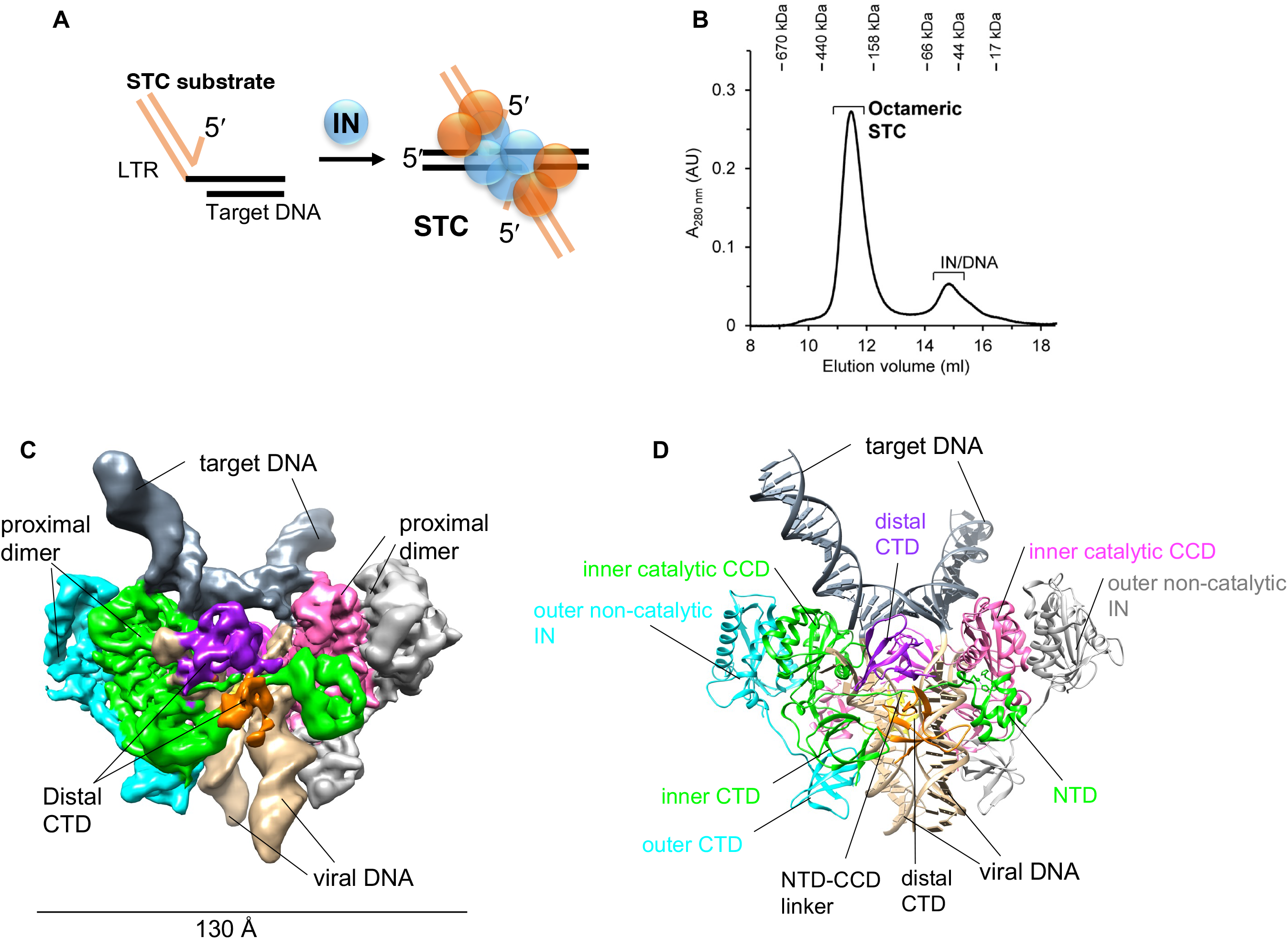
Cryo-EM structure of the RSV strand transfer complex (STC). **A.** Schematics of the STC substrate and RSV IN to assemble STC intasomes. **B.** A typical size-exclusion chromatography (SEC) profile of STC purification. The elution positions of molecular weight standards are marked. **C.** Cryo-EM density of the RSV STC. Each of 8 IN subunits are colored differently. Target and viral DNAs are marked shown in slate gray and tan color, respectively. **D.** Model of the STC structure using same colors as in C. The proximal chains A, B, E, F are shown in pink, gray, green and cyan, respectively while the distal chains C, D, G, H are shown in purple, orange, magenta and yellow, respectively.

**Table 1.** Biochemical characteristics of IN residues described in this study. The experimental details for 3’-processing using a blunt-ended substrate, concerted integration activity, CSC and STC assembly with a recessed 18R substrate are described in the text. The 3’-processing and concerted integration activities and yield of CSC and STC intasome is presented here relative to the wt IN (1-286 IN). “++++”, 75-100%; “+++, 50-75%; “++”, 25-50%; “+”, 10-25 %; “-”, less than 10%.

|  | **3'-processing** | **concerted integration** | **CSC assembly** | **STC assembly** |
| --- | --- | --- | --- | --- |
| wt IN 1-286 | ++++ | ++++ | ++++ | ++++ |
| R244A | ++ | ++ | - | ++ |
| R244E | - | - | - | - |
| Y246A | ++ | ++ | + | +++ |
| S124A | ++++ | ++++ | ++++ | ++++ |
| S124D | ++++ | - | ++++ | - |
| N-C (26 aa) | - | - | - | - |
| C-C (45 aa) | - | - | - | - |
| NC/CC (26/45 aa) | - | - | - | - |

### Interactions between RSV IN and DNA in the STC intasome

IN binding of the target DNA in the CSC intasome stabilizes this complex and produces the STC. We used a computational program DNAproDB (22) to carry out an extensive analysis to identify the potential IN residues that have interactions with DNA in the STC. Most of these interactions were with viral DNA which included contacts with either the nucleotide base only including the major and minor groove (*Fig. S2*) or sugar-phosphate backbone (*Fig. S3*). The majority of the IN interactions were limited to the terminal 10 nucleotides of the non-transferred viral DNA strand.

The NTD-CCD linker of the inner proximal subunit swaps across the CIC in the STC intasome and interacts with viral DNA bound to another unit of proximal dimer in CIC (Fig. 1C, D). The linker residues V50 and P52 interact with T3 and G4 of the non-transferred DNA strand (Fig. 2). Not surprisingly, the proximal inner subunits (shown in green) which provide the active site for catalysis had maximal interactions with the viral DNA nucleotides. At the same time, the outer proximal subunit and distal subunits of IN also had interactions with the viral DNA. There were multiple viral DNA contacts with the distal CTD subunits. For example, the W259, R244 and Y246 (shown in purple) interact with the 5’-terminal nucleotides on the non-transferred DNA strand (Fig. 2).

**Fig. 2.**
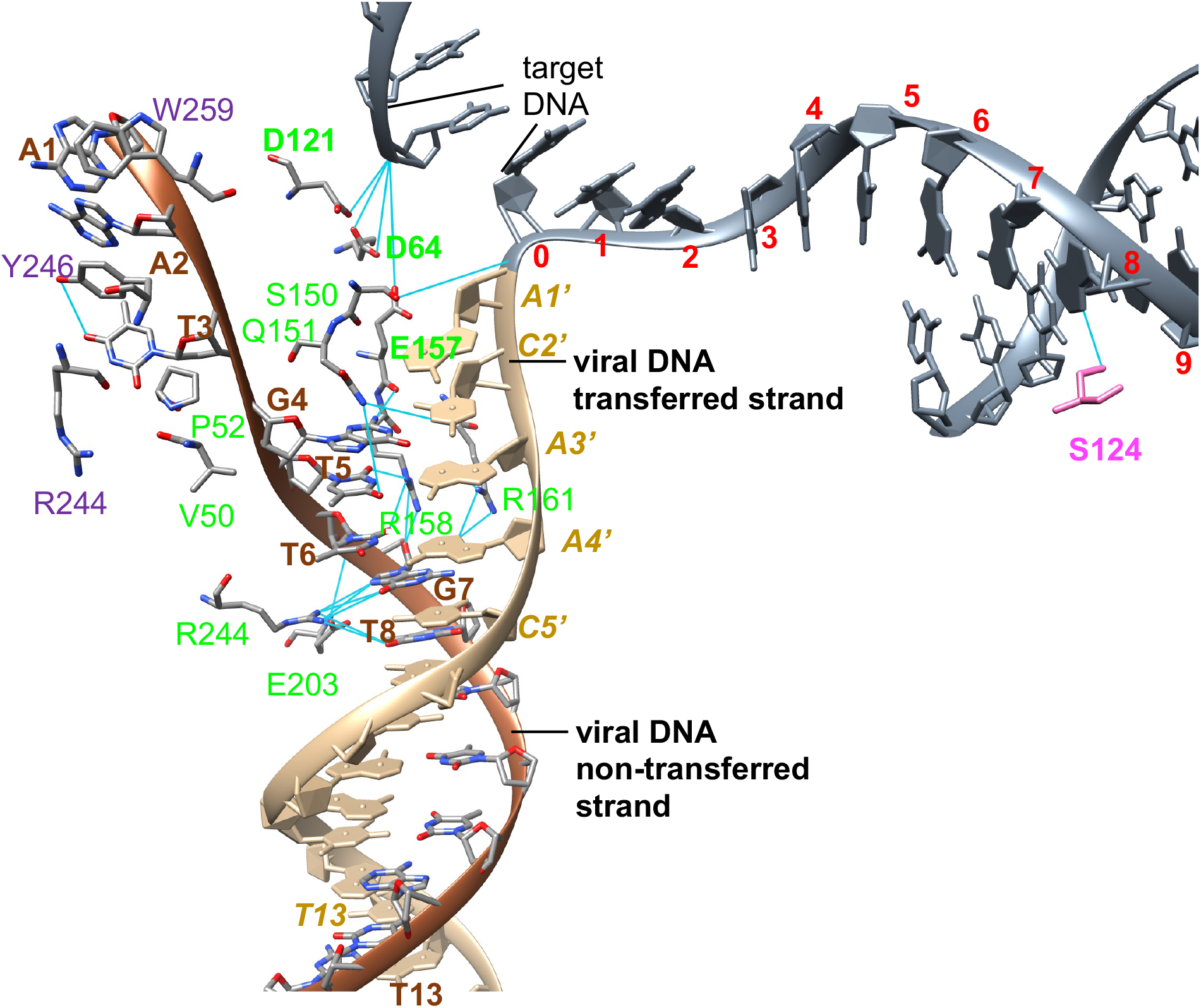
Visualization of selected interactions between IN and DNA in the RSV STC. The two viral DNA strands are depicted in distinct colors. The target DNA (show in slate gray color) is covalently joined to the transferred strand of viral DNA. The select IN residues of proximal and distal subunits interacting with viral DNA nucleotides are colored by their respective protein chain as in Fig. 1D. The catalytic triad of active sites D64, D121 and E157 is also indicated as bold dark green residues.

Compared to the IN contacts with the viral DNA, there were minimal nucleotide base specific contacts in target DNA (Fig. 2, *Fig. S2*). Importantly, S124 of the inner proximal subunits interact with a nucleotide at the 8^th^ position from the cleavage site. However, there were several interactions between IN residues and sugar phosphate backbone of the target DNA, albeit in region beyond the 6 nucleotides after the cleavage site (*Fig. S3*). The 6 nucleotides immediately adjacent to the conserved CA cleavage site had no interactions with IN possibly suggesting the flexibility to accommodate near random integration sites (23).

### Biochemical characterization of IN residues interacting with viral DNA

Based on the cryo-EM structures of the RSV STC and CSC intasome (20), we selected IN residues for site-directed mutagenesis to determine their role in intasome assembly and catalytic activities (Fig. 3 to 5). We expressed and purified RSV IN mutants R244A, R244E and Y246A that interact with the 5’-terminal viral DNA nucleotides of the non-transferred strand (Fig. 2). Similar to wt IN (1–286), all of these IN mutants were dimeric (Fig. S4). We determined the ability of these IN mutants to interact with viral DNA to produce CSC and STC intasomes (Fig. 3). We used longer assembly times and conditions that allow the intermediate tetrameric CSC intasome to be effectively converted into the octameric form by wt IN (Fig. 3A, 3B) (4, 5). Under these conditions, IN mutants R244A and R244E were completely defective in producing tetrameric and octameric CSC intasomes while IN Y246A produced primarily tetrameric CSC intasomes (Fig. 3B). These results suggest that the recruitment of distal dimers to assemble octameric CSC intasomes is compromised with R244A, R244E and Y246A but allow the latter to produce the tetrameric CSC. IN mutant R244E was unable to assemble STC while R244A and Y246A were diminished in their ability to produce STC compared to the wt IN (1-286) (Fig. 3C). These intasome assembly results matched with the anticipated catalytic activities of these IN mutants. The catalytic activities were determined by measuring the 3’-processing and strand transfer activities to produce concerted integration products (Fig. 5, Table 1). Using blunt-ended viral DNA substrates, the 3’-processing assays were performed in presence of Mg^++^ as the divalent metal ion needed for catalysis. The wt IN (1-286) showed ∼35% of 3’-processing activity (Fig. 5A, Table 1). The mutant R244A had reduced 3’-processing activity using blunt-ended substrate as well as reduced concerted integration activity determined with recessed 18R substrate (Fig. 5B). R244E was devoid of 3’-processing and concerted integration activity. Y246A was partially active (∼50% of wt IN) for 3’-processing (Fig. 5A) and for concerted integration using a 3’-OH recessed substrate (Fig. 5B). As mentioned, Y246A is also defective in assembly of octameric CSC intasome suggesting it is deficient in recruiting the distal subunits. All of these IN mutants were defective to various degrees for assembling the STC intasome suggesting multiple functions associated with these residues, in contrast to other IN single-point mutants that do not affect assembly of the STC intasome (5).

**Fig. 3.**
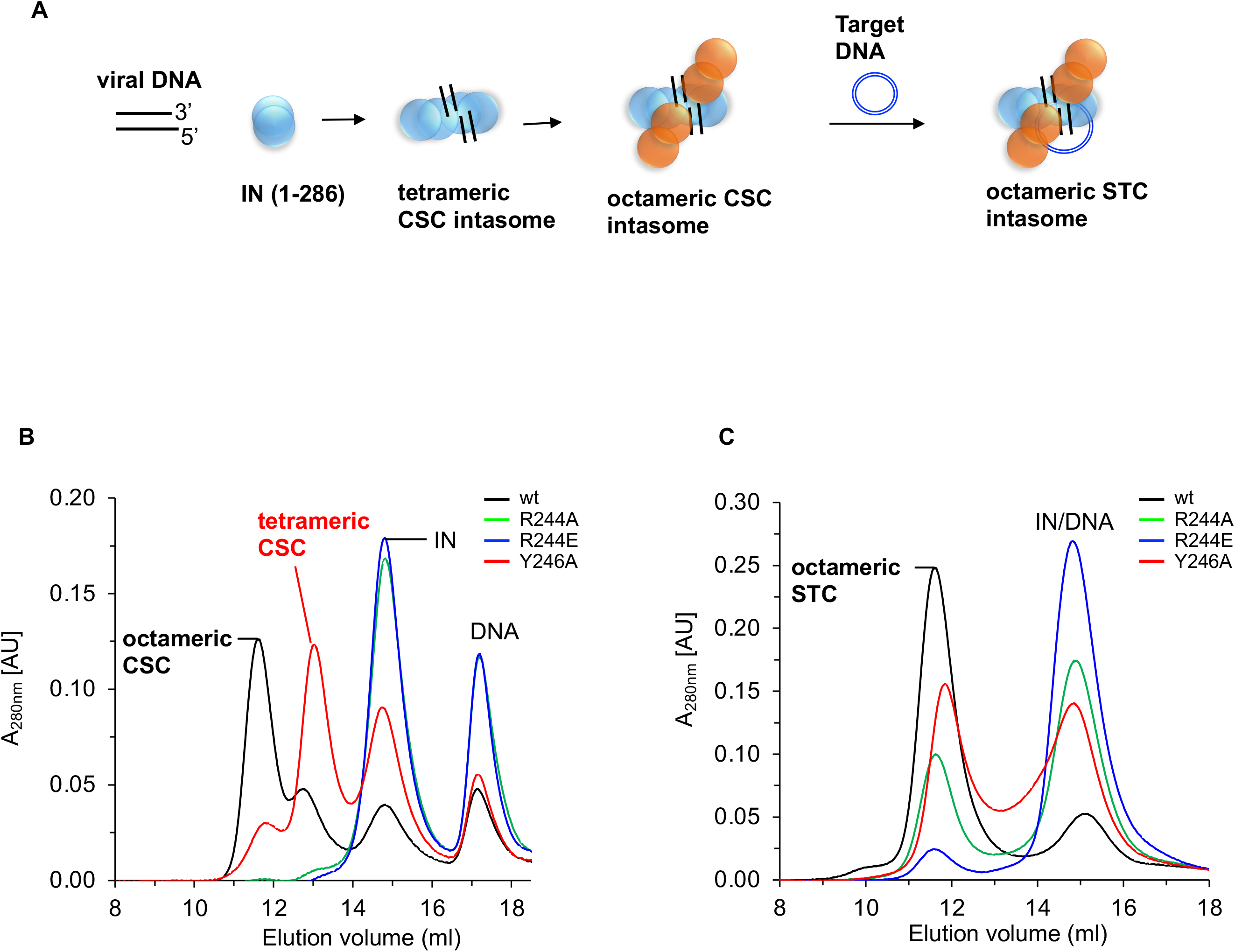
Effects of substitutions at select IN residues interacting with viral DNA for CSC and STC intasome assemblies. **A.** Schematics of RSV intasome assembly. The wt full-length IN (1-286 aa) binds to viral LTR DNA ends, resulting in the assembly of a transient intermediate-tetrameric CSC intasome. IN is shown as dimer. Binding of additional IN molecules in distal positions results in the formation of a stable octameric CSC. In presence of a target DNA, an octameric strand transfer complex (STC) is formed. Proximal and distal IN protomers are shown in blue and orange, respectively**. B.** MK-2048 trapped CSC intasomes were assembled with wt RSV IN or its mutants R244A, R244E and Y246A at 18°C for 18 h. The assembled CSC intasomes were analyzed by SEC. **B.** The STC intasomes were assembled for 18 h at 4°C with the above IN proteins and analyzed by SEC. Elution positions of molecular weight markers are indicated. AU, arbitrary units.

**Fig. 4.**
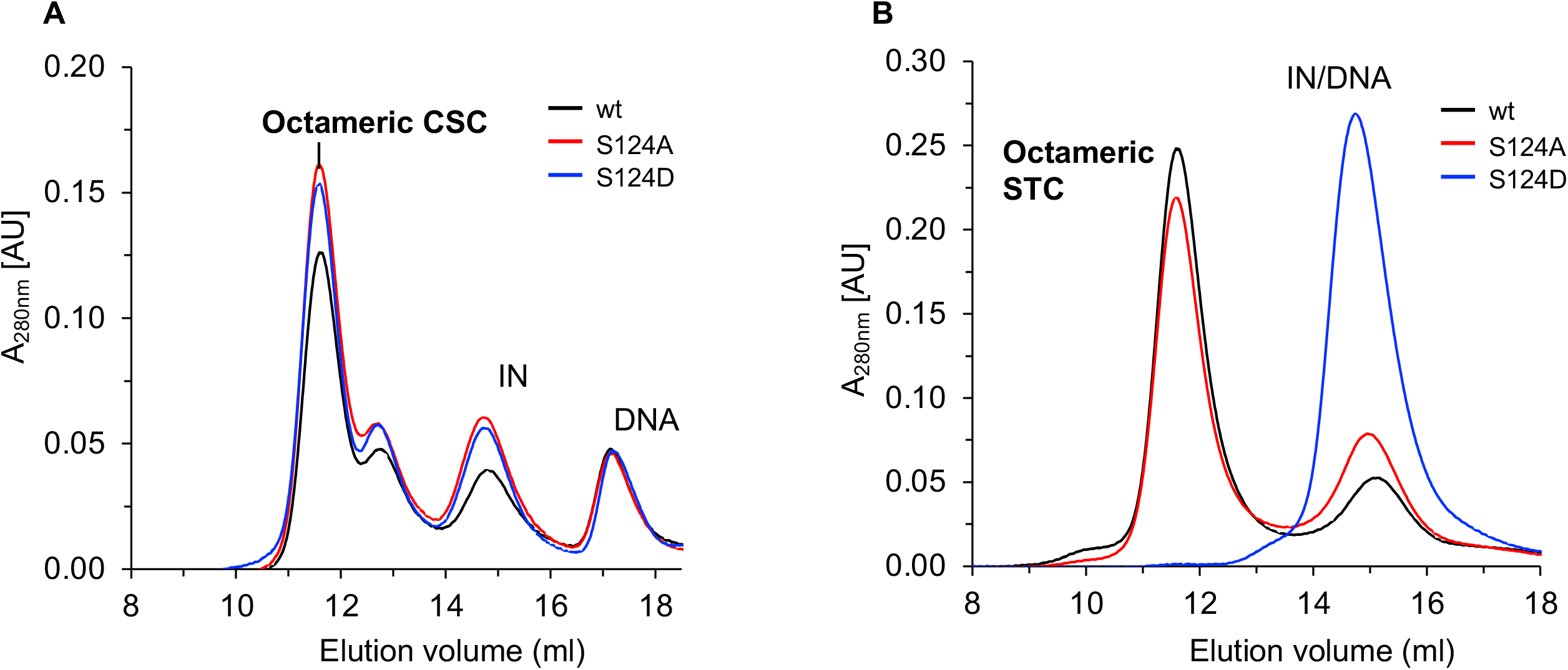
Analysis of S124 substitutions on CSC and STC assemblies. **A.** MK-2048 trapped CSC intasomes were assembled with wt RSV IN (1-286) and IN mutants S124A or S124D at 18°C for 18 h. The assembled CSC intasomes were analyzed by SEC. **B.** The STC intasomes were assembled for 18 h at 4°C with these same IN proteins and analyzed by SEC. Elution positions of molecular weight markers are indicated. AU, arbitrary units.

**Fig. 5.**
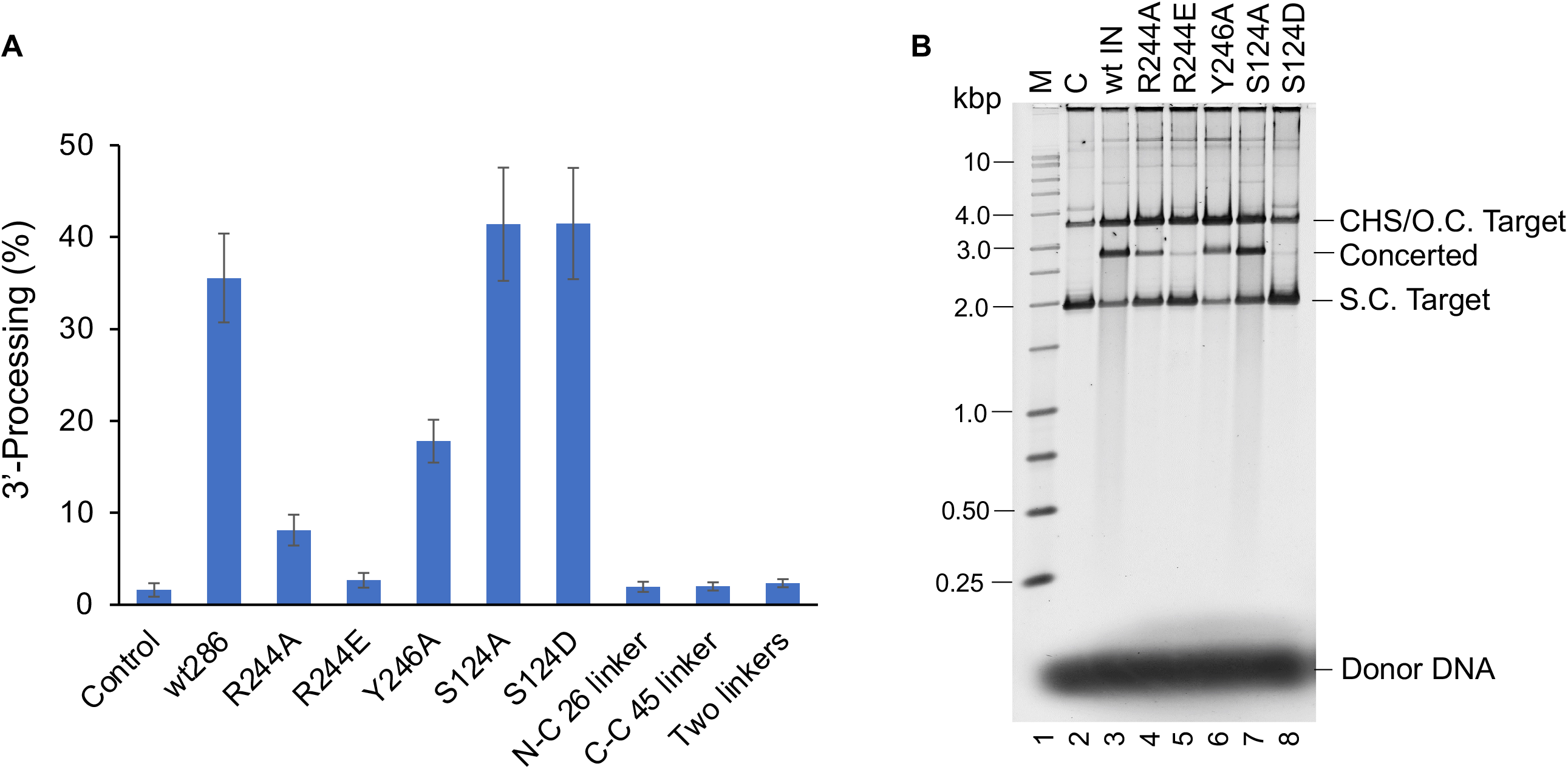
Catalytic activities of RSV wt and mutant IN. **A.** 3’-processing activities of wt IN (1-286) and designated mutant IN proteins were determined using 20 nM IN and 0.5 nM of ^32^P labeled blunt-ended DNA substrates at 37°C for 30 min. The standard deviation was determined from at least three independent experiments. **B.** The concerted activity of each IN mutant was determined with 18R GU3 substrate at 37°C for 30 min. The products were deproteinized and run on a 1.3% agarose gel. Lane 1, marked M contains the molecular size marker (Promega kb ladder). Lane 2, marked C does not contain IN. lanes 3-9 contain different IN as indicated on the top. CHS, circular half-site; s.c. supercoiled; o.c. open circular. Unused donor DNA is indicated at bottom right.

### Interactions of RSV IN residues with target DNA

We observed minimal nucleotide specific interactions between IN and target DNA (Fig. 1D, *Fig. S2*). Previous data suggested that RSV IN S124 played a direct role in target DNA binding (24). We determined that S124 of inner catalytic subunit interacts in the minor grove with the third nucleotide adenine from the 6 nucleotide duplication sequence. In addition, S124 has number of interaction with target DNA backbone with sugar and phosphate groups. These interactions of S124 with the target DNA backbone seem to be position specific rather than the nucleotide specific. We made substitution at S124 to a neutral amino acid alanine or to a negatively charged residue aspartic acid.

As expected, these substitutions did not affect CSC intasome assembly since S124 interactions were limited to target DNA only (Fig. 4A). IN mutant S124A showed similar efficiency to assemble STC while S124D was completely deficient in producing the STC (Fig. 4B). Correspondingly, the IN mutant S124A had similar 3’-processing and concerted integration activity comparable to wt IN (Fig. 5A). In sharp contrast, S124D had 3’-processing activity similar to wt IN but was devoid of concerted integration activity (Fig. 5A and 5B). The results demonstrate that S124 is essential for binding to target DNA to assemble the octameric STC.

### Role of inter-domain linker regions in RSV intasome assembly

It has been speculated that length of the inter-domain linker may determine the oligomeric form of retroviral intasomes. PFV IN has relatively longer inter-domain linkers compared to RSV, mouse mammary tumor virus (MMTV) and HIV-1. To determine the role of linker length in RSV IN, we modified the NTD-CCD as well as CCD-CTD linker lengths (13 and 8 aa respectively) in RSV IN to mimic PFV IN either individually or together. WT RSV IN (1-286) constructs possessing a 26 aa NTD-CCD linker (N-C 26) or a 45 aa CCD-CTD linker (C-C 45) or both of these linkers (NC/CC) were constructed. All linker modified INs were dimeric (*Fig. S4*). Surprisingly, all three linker modified IN constructs were inactive for 3’-processing (Fig. 5A) as well as concerted integration (data not shown). The alterations in inter-domain linker lengths also abolished the CSC and STC intasome assemblies. These data suggest that inter-domain linkers in different retroviral INs have evolved naturally and a change in their length abolishes the intasome assembly and catalytic activities.

## Discussion

In the RSV concerted integration pathway, IN binds to the viral DNA ends producing the intermediate tetrameric CSC intasome followed by the formation of the mature octameric CSC. The mature octameric CSC binds target DNA followed by strand transfer producing the STC intasome. We assembled the STC intasome that contains IN bound with a covalently linked viral/target DNA substrate. In this study, we determined the structure of RSV STC intasome by single-particle cryo-EM and investigated structure-function relationships between the CSC and STC intasomes. We previously determined the structure of octameric CSC stabilized by the INSTI MK-2048 by single particle cryo-EM (20). Three-dimensional variability analysis of the RSV STC structure demonstrated significant flexibility of the distal subunits similar to the RSV octameric CSC intasome produced with IN and viral DNA only (20). The conformational flexibility of distal subunits specifically in the NTD-CCD regions in the STC was responsible for insufficient electron density coverage (*Fig. S1D*). We speculated that this flexibility of distal subunits is required to accommodate random target DNA sequences for integration.

We previously determined the structure of RSV octameric STC by x-ray crystallography (15). The crystal structure contained IN (1-270) which possessed a multitude of single-point substitutions C23S, L112M, L135M, L162M, L163M, L188M and L189M. In crystal form, the STC is locked in a single conformation via crystal lattice packing. In contrast, wt IN (1-278) was used to assemble the RSV octameric STC for structure determination by cryo-EM. These IN-DNA complexes are in-solution and hence free to adopt a wide variety of conformations. The cryo-EM structure of STC showed improved overall resolution (3.36 Å) and nearly 3 Å in the CIC region as compared to the crystal structure (3.86 Å). The CIC region of the STC obtained by cryo-EM was similar to CSC intasome (20) and the crystal structure of STC (15). Overall, the root mean square deviation (RMSD) of C-*α* atoms in IN subunits was 0.805-1.1508 (*Fig. S5*) between the cryo-EM structure of RSV STC and its crystal structure (15) across 856 residues. However, the NTD-CCD of the distal subunits remained flexible and poorly resolved in the cryo-EM structure of STC. In the crystal structure of the STC, the CCD of distal subunits were loosely associated with distal region of target DNA through non-specific interactions. Similar to the cryo-EM structure of STC, there were no interactions identified beyond the 11^th^ residue on target DNA from the viral-target DNA junction. Most of the interactions in the target DNA are limited to the CCD region of proximal IN subunits which also contributes the active site (Fig. 2, *Fig. S2*). Absence of IN interactions in outlying target DNA region possibly contributes to the flexibility observed in the distal NTD-CCD regions in STC structure obtained by cryo-EM (*Fig. S1D, S6*).

We identified several potential IN-DNA interactions critical for CSC and STC intasome assemblies. Selected interactions (R244, Y246 and S124) were investigated for structure-function relationship studies using site-directed mutagenesis to determine their role in CSC and STC intasome assembly and catalytic activities. R244 of the distal IN subunit interacts with T3 at the 5’ terminal non-transferred strand in viral DNA suggesting a critical binding point for distal IN subunits (Fig. 2). The proximal subunits interact with viral DNA at nucleotides 6-8 on non-transferred strand (Fig. 2, *Fig. S2*). We made substitution mutant R244A and R244E to establish the distal subunit DNA binding properties. Both the R244A and R244E IN substitutions were unable to assemble octameric CSC intasome (Fig. 3B) but could assemble the STC at a reduced capacity (Fig. 3C). It’s probable that target DNA helped stabilize the STC intasome assembly by providing additional interaction sites. RSV IN (1-269) which produces a tetrameric CSC only is able to assemble octameric STC in vitro (4). The G7 nucleotide which interacts with R244 has been shown to be critical for concerted integration (25). It seems likely that R244 has multiple roles; *i.e.* in binding to the viral DNA via proximal subunits as well in the CIC region assembly through interactions with CTD of distal subunits. The corresponding residue E246 in HIV-1 was shown to interact with viral DNA (26) and later studies study determined that substitution at E246 to E246A/K affect 3’-processing and concerted integration (27,28). RSV IN W259 residue interacts with terminal nucleotide on non-transferred strand (Figs. 2, *S2*) and has been shown to affect the CTD-CTD interaction of distal IN subunits. Substitutions of W259 to A/R/T abolished the 3’-processing and strand transfer activities (29).

The first and third nucleotide (adenine and thymine, respectively) from the 5’-end of the non-transferred end interacts with Y246 of the CTD from a distal subunit (Fig. 2). Y246A IN assembled only the tetrameric CSC intasome with significantly reduced quantities of mature octameric CSC intasome indicating its critical role in octameric CSC assembly (Fig. 3B). Whether this tetrameric CSC intasome containing two proximal dimers reflects the transient intermediate (4, 5) in concerted integration pathway is unclear and warrants further investigation. Y246A showed moderate efficiency to assemble STC intasome (Fig. 3C) and possessed deceased 3’-processing and concerted integration activity (Fig. 5). There appears to be no other apparent interactions of Y246 with target DNA in the STC intasome (*Fig. S2*). In addition, the possibly exists that 3’-OH processing occurs in the tetrameric intasome (Fig. 3B) because Y246 binds to T3 (distal subunit) and T5 (inner catalytic subunit of proximal dimer) of the 5’ end of the non-transferred DNA strand in the octameric intasome (*Fig. S7*) (20) which may be partially responsible for distortion of DNA blunt-ends necessary for 3’-OH processing (30). In summary, it seems possible that the disruption of Y246 of the distal subunit interaction with the non-transferred viral DNA strand (Fig. 2) affects the recruitment of the distal subunits for octameric intasome assembly.

The CSC intasome assembly and its conversion to STC does not induce drastic conformational changes. Similar to RSV, earlier studies of PFV (13, 31) and MVV (16) intasomes, which are tetrameric and hexadecameric intasomes, respectively, also showed essentially no changes before and after target DNA binding. IN is well positioned to bind the target DNA in RSV octameric CSC intasome. The root mean square deviation (RMSD) of C-*α* atoms in IN subunits was 0.668-1.132 (*Fig. S8*) between the cryo-EM structures of RSV CSC (20) and STC structures across 917 residues. There were minimal interactions between IN and six nucleotides immediately downstream of viral and target DNA junction. The crucial backbone specific interactions are between S124 and 8^th^/9^th^ nucleotide downstream to the viral-target DNA junction site (*Fig. S2, S3*). A similar IN residue-nucleotide position interaction in the STC is conserved across multiple retrovirus species (32). The S124 residue in RSV IN was previously implicated in target site selection and replication in vivo (24,32,33). The S124A substitution has no drastic effects on the CSC or STC intasome assembly (Fig. 4). As expected, S124A had 3’-processing and concerted integration activities similar to the wt IN (Fig. 5). These results are in full agreement with earlier studies including the fact that RSV virions with the S124A IN mutation replicates similarly to the wt RSV (24). However, RSV containing IN with the S124D substitution was unable to replicate in cell culture (33). An S124D substitution is unique that it separated the two biologically relevant enzymatic activities of IN. It allows the 3’-processing of the viral DNA substrates but blocks the strand transfer into host DNA. Our studies show that the S124D substitution had no effect on CSC intasome assembly (Fig. 4A) but STC intasome assembly was completely inhibited (Fig. 4B). Likewise, no strand transfer activities were detected (Fig. 5B) thus providing direct evidence that binding to the target DNA is affected. In summary, our studies of S124 provide evidence for its role in RSV CSC and STC intasome assemblies and structural rationale into its effect on catalysis and viral replication.

While this work was in progress a study by Jozwik et al. (34), reported similar IN-viral and target DNA interactions in the MMTV STC intasome. Collectively, our data advance the understanding of RSV intasome structure and function and identified molecular determinants involved in intasome assembly.

## Material and Methods

### RSV IN expression and purification

RSV IN constructs (wt, C-terminal tail truncations and single-point mutants) were expressed in *Escherichia coli* BL21 (DE3) pLysS and purified to near-homogeneity (3,5,35). The wt Prague A IN subunit is 286 aa in length, designated 1-286. Purified IN were concentrated to 20-30 mg/ml using Amicon Ultra-15 (30K MWCO) centrifugal filters. IN mutant expression constructs (R244A, R244E, Y246A, S124A and S124E were produced in full length RSV IN (1-286) by site-directed mutagenesis. The NTD-CCD and CCD-CTD linker region of RSV IN (1-286) were modified to mimic the corresponding regions from PFV IN (13) either individually or together. The modified construct having the 26 aa NTD-CCD linker and the 45 aa CCD-CTD linker were named N-C 26 and C-C 45, respectively. The IN construct possessing both linkers was termed NC/CC (26/45). The DNA sequence of all IN constructs was confirmed by sequencing. The protein concentrations were expressed as monomeric subunits.

### Assembly and purification of the RSV STC intasome

The RSV octameric STC intasome was assembled with IN 1-278 and STC DNA substrate (4, 15). The STC substrate was prepared by annealing three oligonucleotides in equimolar ratio (42 nt 5’-GAGTATTGCATAAGACAACAGTGCACGAAAGAAGAAGACACT-3’, 22 nt 5’-AATGTTGTCTTATGCAATACTC-3’ and 16 nt target 5’-AGTGTCTTCTTCTTTC-3’. The IN and STC substrate (35 μM and 10 μM, respectively) were mixed in 20 mM HEPES, 1 M NaCl, 20% glycerol, 1 mM TCEP pH 7.5 and dialyzed in a Slide-A-Lyzer G2 cassette (3.5 K cut off) overnight against 20 mM HEPES, 0.125 M NaCl, 20% glycerol, and 1 mM tris(2-carboxyethyl phosphine)(TCEP), pH 7.5. Precipitated intasomes were solubilized by dialysis against a higher salt buffer 20 mM HEPES, 0.75 M NaCl, 20% glycerol, and 1 mM TCEP, pH 7.5 for 1 h at room temperature. The octameric RSV STC intasomes were purified by SEC using Superdex 200 Increase column (10/300)(Cytiva Life Sciences) (Fig. 1B). The SEC running buffer was 20 mM HEPES, pH 7.5, 650 mM NaCl, and 1 mM TCEP. SEC purified fractions were used immediately for vitrification. Re-chromatography of pooled STC fractions on Superdex 200 established that the octameric RSV STC intasomes were stable on ice even after overnight storage at 4°C.

### Concerted integration and 3’-OH processing assays

The concerted integration assays were performed using 3’ OH recessed oligonucleotide viral DNA substrates with RSV IN as previously described (3, 4). Double stranded 3’ OH recessed substrates containing RSV gain-of-function(G) U3 and wt U3 long terminal repeat (LTR) sequences were 18 nucleotides in length and synthesized by Integrated DNA Technologies. The DNA substrates were recessed by two nucleotides on the catalytic strand and designated with an R. The identified length of the oligonucleotide denotes the non-catalytic strand. The sequences were as follows: GU3 18R (5’-ATTGCATAAGAC**<u>A</u>**ACA-3’ and 5’-AATGT**<u>T</u>**GTCTTATGCAAT-3’). The bold underlined nucleotide on the catalytic strand were different between the GU3 and wt U3 sequence. The concentrations of IN and the viral ODNs in a typical assay were 2 and 1 µM, respectively. The strand transfer products were separated on a 1.3% agarose gel, stained with SYBR Gold (Invitrogen) and analyzed by a Typhoon 9500 Laser Scanner (GE Healthcare Life Sciences).

The 3’-OH processing of ^32^P-labeled blunt-ended viral 4.6 kb DNA at 37°C was previously described (29, 36). Concentrations of purified IN and DNA in the assay mixture was 20 nM and 0.5 nM, respectively.

### Assembly of the RSV CSC

To determine the effect of IN mutations, the RSV CSC was assembled with IN and GU3 18R in the presence of MK-2048 (3). The assembly buffer was 20 mM HEPES, pH 7.5, 100 mM NaCl, 100 mM ammonium sulfate, 1 M non-detergent sulfobetaines (NDSB)-201, 10% dimethyl sulfoxide (DMSO), 10% glycerol, 1 mM TCEP. IN (as monomers), 3’ OH recessed DNA ODN and MK-2048 concentrations were 45 µM, 15 µM and 125 µM, respectively. After addition of MK-2048 and DNA, IN was added to the assembly mixture. MK-2048 was generously provided by Merck & Co. The samples were generally incubated at 18°C for 18 h. The octameric RSV CSC intasomes were analyzed by SEC using Superdex 200 Increase (10/300). The SEC running buffer was 20 mM HEPES pH 7.5, 200 mM NaCl, 100 mM ammonium sulfate and 1 mM TCEP. MK-2048 was omitted from the running buffer. Chromatography was performed at 4°C and UV absorption monitored at 280 nm.

### Cryo-EM sample preparation of the RSV STC and imaging

The **c**ryo-EM samples derived from the peak SEC fractions (∼0.4 µM) were used promptly for vitrification. The samples were prepared on quantifoil holey carbon grids (R2/2 300 mesh copper), which were plasma cleaned for 1 min using a Gatan Solarus 950 (Gatan) and plunge frozen using a Vitrobot Mark IV (ThermoFisher Scientific). The Vitrobot sample chamber was set to 4°C and 100% humidity. 3 µl of SEC purified RSV STC was applied to the plasma cleaned quantifoil grids and allowed to incubate for 20 sec. Samples were then blotted for 2 sec at a blot force of -1 and plunge frozen into liquid ethane. Vitrified grids were imaged using a Cs-corrected ThermoFisher Titan Krios G3 electron microscope (ThermoFisher Scientific) operating at an accelerating voltage of 300 kV equipped with a Falcon 4 detector (ThermoFisher Scientific). Data acquisition was automated using EPU software (ThermoFisher Scientific) at a magnification of 59,000x which corresponds to a pixel size of 1.16 Å. Movies were recorded for 13.29 sec with 50 frames with a dose rate of 1.0 electrons per Å^2^ per frame (total dose of 50 electrons per Å^2^). The defocus was varied between -1 to -2.5 µm. A total of 3297 movies were recorded including 248 movies at a 20-degree stage tilt. The data collection parameters are indicated in *Table S1*.

### Cryo-EM data processing

The movies were corrected for beam induced movement using MotionCorr2 (37). Further data analysis was done in cryoSPARC 3.0 (*Fig. S1*) (21). The contrast transfer function (CTF) was determined using Patch based CTF estimation. The micrographs with estimated CTF fit resolution in range of 2.4 to 6.0 Å were selected for further analysis. Initially, the blob picker was used to particles of diameter 120-200 Å from 500 micrographs. The particles were extracted using a box size of 324 pixel and reference free 2-D classification performed. The selected 2D classes showing different conformations were used as a template to pick particles from all micrographs. The 867, 209 particles were subjected to 2D classification in 100 classes. After removing the junk particles, select 2D class-averages containing 214, 690 particles were used for Ab-initio 3D reconstruction. The 3D reconstructions were refined by homogeneous refinement without imposing symmetry. The class with clear feature showed a global resolution map of 3.98 Å at Fourier shell correlation of 0.143. The non-uniform refinement of this class using C2 symmetry resulted in a 3.36 Å global resolution map and was used for model building and refinement. Heat map of angular distribution of refined particles used in reconstruction is shown in (*Fig. S1G*). Directional 3DFSC curves were calculated using the wrapper program within the cryoSPARC (38). The local resolution in the CIC region was determined in cryoSPARC and displayed in Chimera (*Fig. S1*). The local resolution map demonstrated ∼2.8-4.0 Å resolution in the core region.

### Preparation of the atomic model, refinement and validation

Our previously reported RSV STC crystal structure at 3.86 Å resolution (PDB ID 5EJK) was used as the preliminary model to dock into the EM map as a rigid body and manually modified/rebuilt using COOT (39). The preliminary model was refined using PHENIX (40, 41) against the cryo-EM density and a standard set of geometry/stereochemistry restraints.

## Funding and additional information

This work was supported by the National Institutes of Health (NIAID AI165081 to KKP, and NIGMS GM118047 to HA). This work was also supported by Saint Louis University President Research Fund (KKP), and Core Usage Fund (KKP) provided by the Washington University Institute of Clinical and Translational Sciences which is, in part, supported by the NIH/National Center for Advancing Translational Sciences (NCATS), CTSA grant #UL1 TR002345. The content is solely the responsibility of the authors and does not necessarily represent the official views of the National Institutes of Health.

## Data availability

The cryo-EM maps were deposited with the Electron Microscopy Data Bank (accession code EMDB-27823) and the refined model with the Protein Data Bank (8E14). All materials used in the manuscript are available upon request.

## Supporting information

This article contains Supporting Information (Figs. S1-S8).

## Acknowledgements

We would like to thank Michael Rau and Dr. Brock Summers for their help in data collection on Krios at Washington University Center for Cellular Imaging.

## Abbreviations

IN: integrase
RSV: Rous sarcoma virus
HIV-1: human immunodeficiency virus type-1
PFV: prototype foamy virus
MMTV: mouse mammary tumor virus
MVV: maedi-visna virus
CSC: cleaved synaptic complex
STC: strand transfer complex
CIC: conserved intasome core
NTD: N-terminal domain
CCD: catalytic core domain
CTD: C-terminal domain
LTR: long terminal repeat
STI: strand transfer inhibitor
wt: wild type
aa: amino acids
SEC: size exclusion chromatography
CHS: circular half site
MW: molecular weight
AU: arbitrary units

## Supporting information

**Fig S1.**
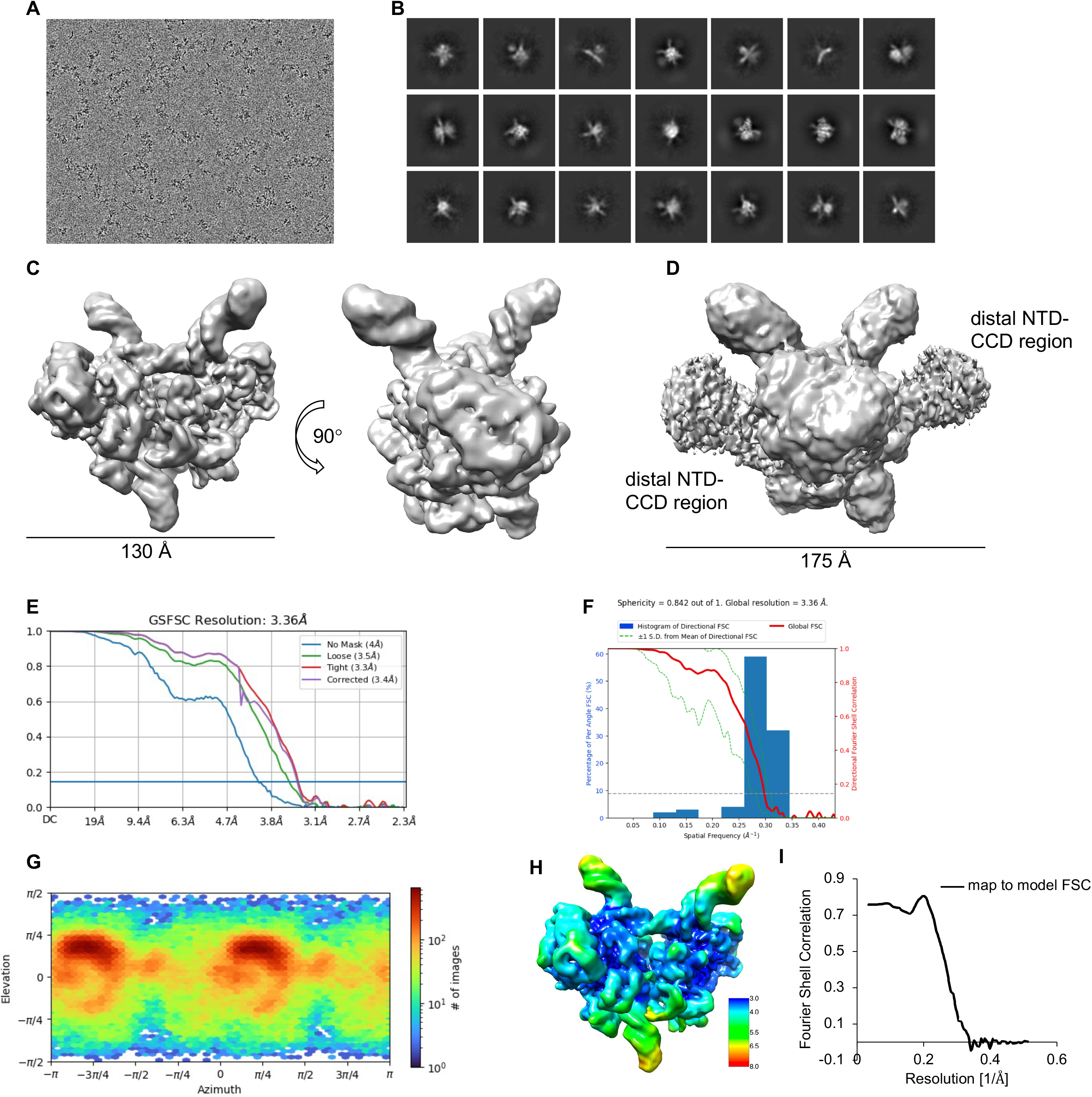
Cryo-EM data analysis workflow. **A.** Representative cryo-EM micrograph. **B.** Representative reference free 2-D class averages of the particles after template picking. **C.** Non-uniform refinement of the 3D map using C2 symmetry. **D.** Same map as in C shown with lower threshold to show flexible NTD-CCD regions of distal dimer subunits. **E.** Gold standard Fourier shell correlation curves (FSC) for the half-maps. **F.** Global resolution FSC curve overlaid onto a histogram of direction resolution value. **G.** Particle viewing directional distribution plot of the RSV STC. **H.** Reconstruction of STC map colored by local resolution. **I.** map to model FSC plot. Detailed description of data analysis is provided in Materials and Methods.

**Fig S2.**
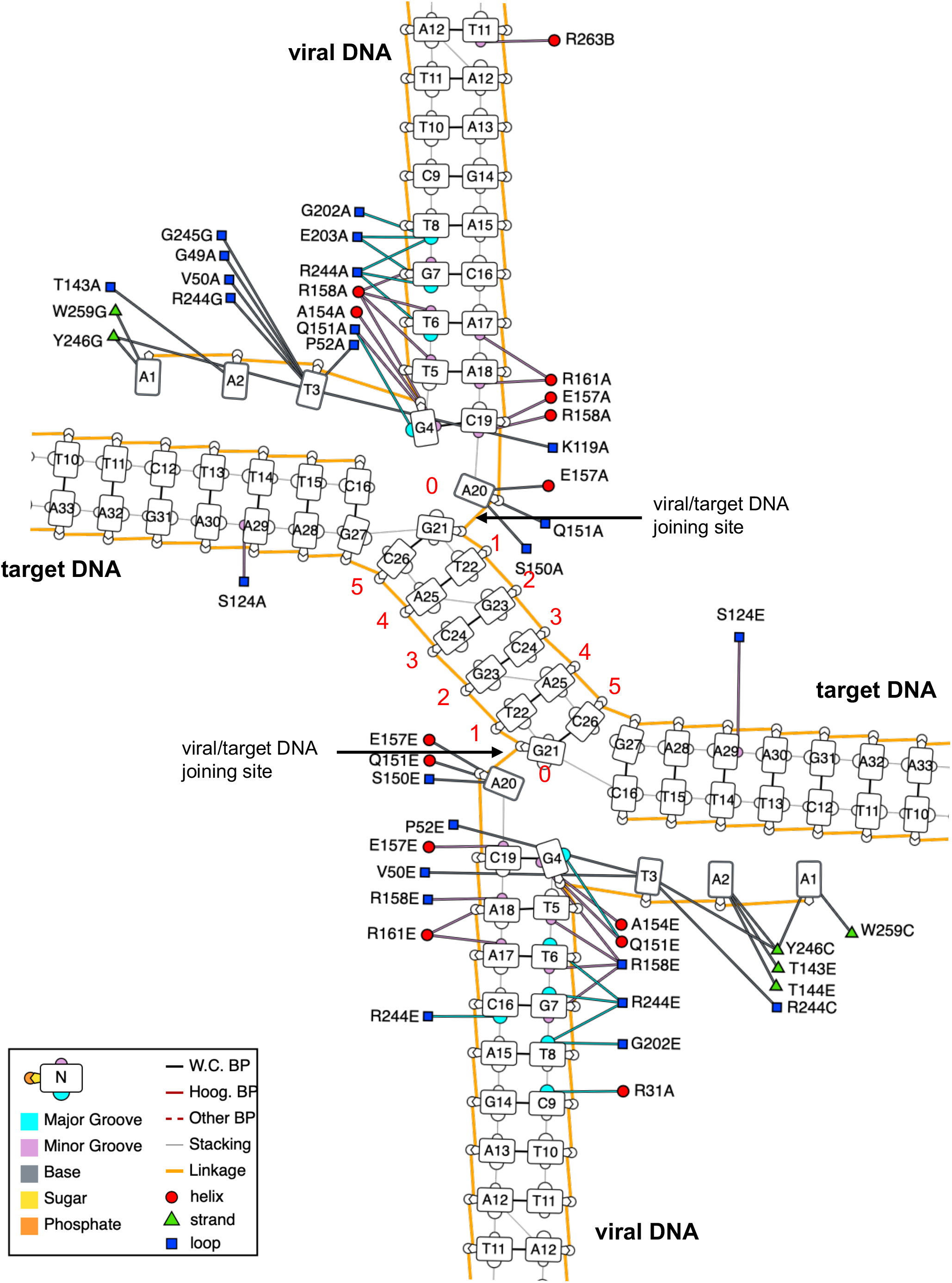
Potential nucleotide base-specific interactions between IN and DNA in the RSV STC. Analyses were performed using DNAproDB to determine nucleotide base specific interactions including major and minor grove. Most of the IN interactions are with viral DNA and minimal interaction with target DNA occurs. Protein residues are labeled with their one letter codes, respective residue number and the protein chain identifier (*e.g.*, R244E indicate R244 of chain E). The protein chains follow the same designation as in Fig. 1D. The 6 bp host site duplication sequence is labeled 0-5 in red. The site joining the viral DNA to target DNA is marked. Schematic shows the potential interactions between IN subunits and nucleotide bases, major and minor grooves. W.C. BP – Watson–Crick bp; Hoog. BP – Hoogsteen bp; Other BP – other bp.

**Fig S3.**
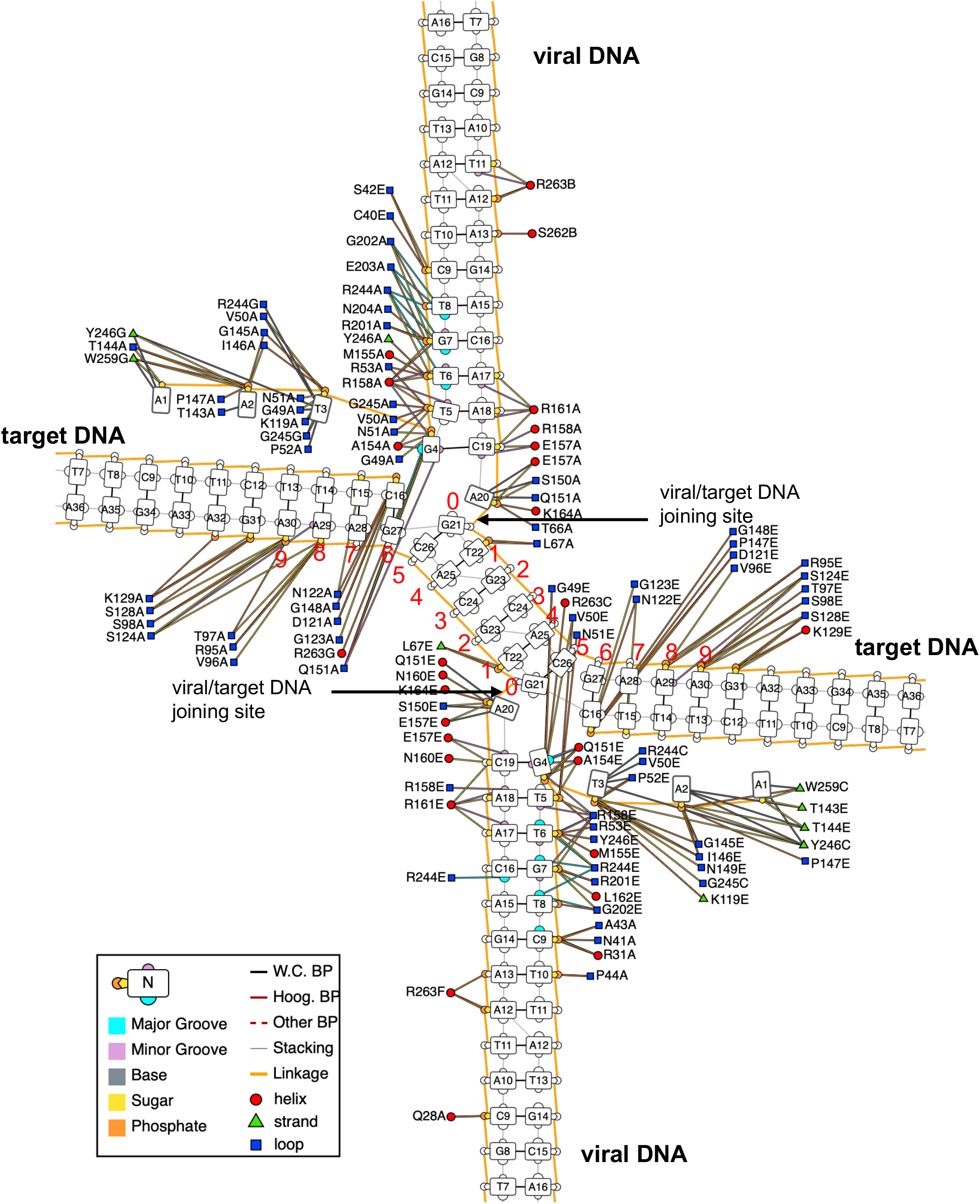
Potential backbone interactions between IN and DNA in the RSV STC. The analyses were performed using DNAproDB. There were minimal interaction in the six bp duplication region at the viral-target DNA junction. The majority of the IN interactions with target DNA are in region beyond the 6 bp duplication region. Protein residues are labeled with their one letter codes, respective residue number, and the protein chain identifier (*e.g.*, R244E indicate R244 of chain E). The protein chains follow similar designation as in Fig. 1D. The 6 bp host site duplication sequence is labeled 0-5. The site joining the viral DNA to target DNA is marked. Schematic shows the interactions between IN protomers and nucleotide bases, major and minor grooves. W.C. BP – Watson–Crick bp; Hoog. BP – Hoogsteen bp; Other BP – other bp.

**Fig S4.**
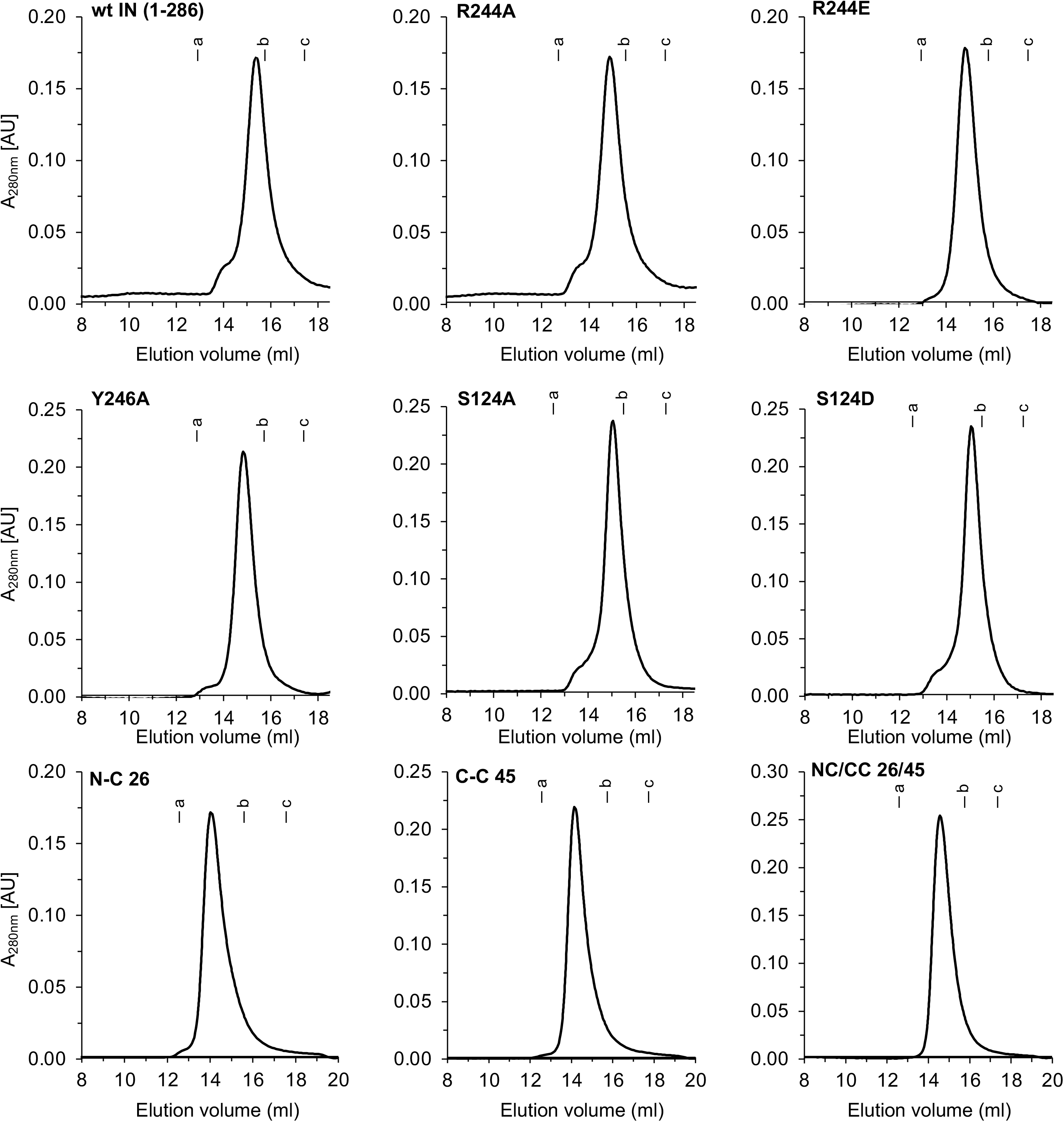
Oligomeric forms of RSV IN mutants. wt RSV IN (1-286) and its indicated variants containing missense substitutions and alterations in the inter-domain linker length were analyzed by SEC using Superdex 200 Increase column (10 x 300 mm). All of the IN proteins at 45 μM concentrations were incubated in the CSC intasome assembly buffer overnight at 4°C before SEC analysis. IN eluted predominantly as dimeric species. The molecular weight standards were run in parallel with each analysis and their elution positions are marked on top of each profile. The slight variation in relative elution positions among different SEC run was due to use of a new column for few samples.

**Fig. S5.**
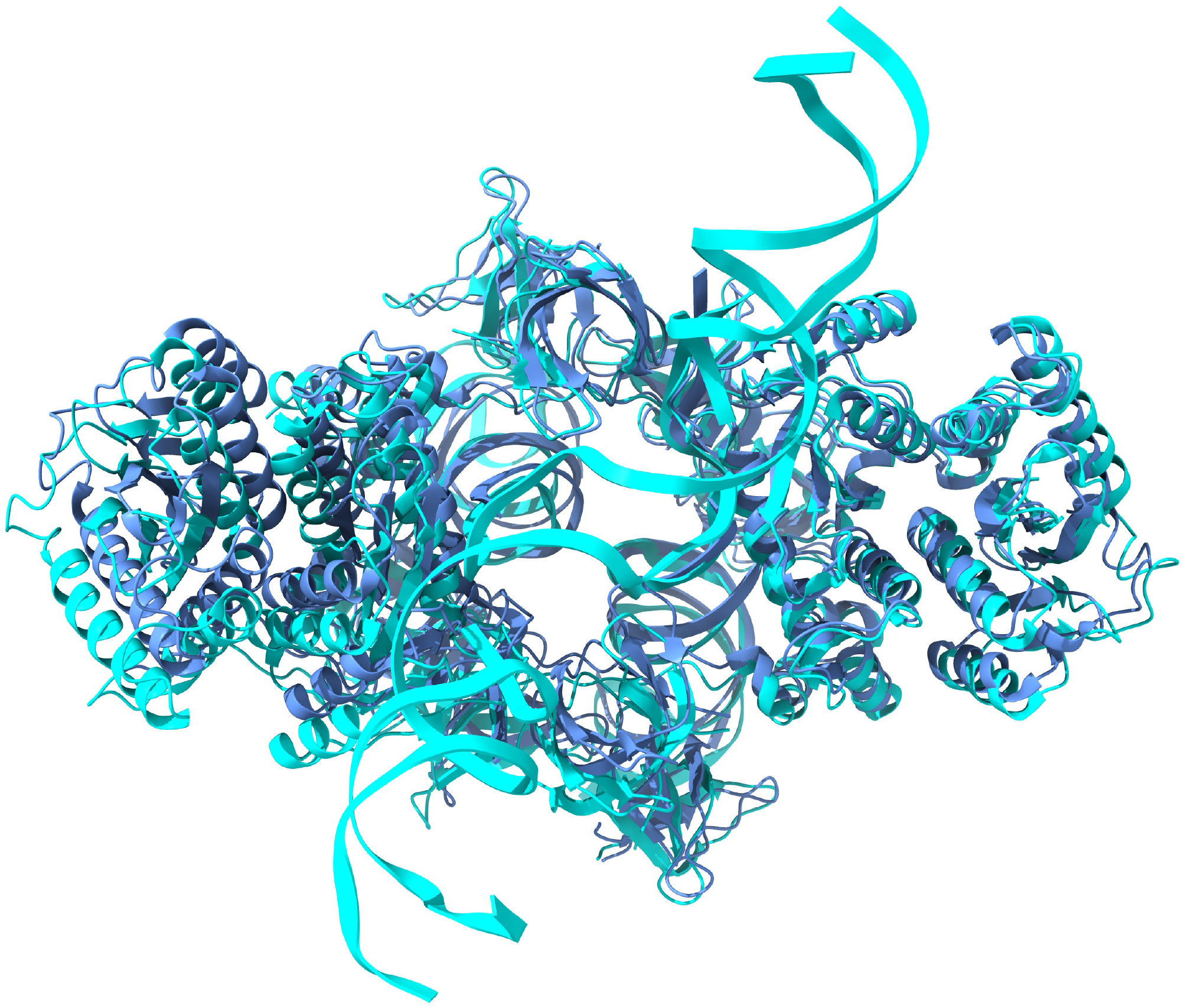
Comparison of the CIC region in RSV CSC intasome (PDB 7JN3, shown in blue) and STC (in cyan) determine by cryo-EM.

**Fig. S6.**
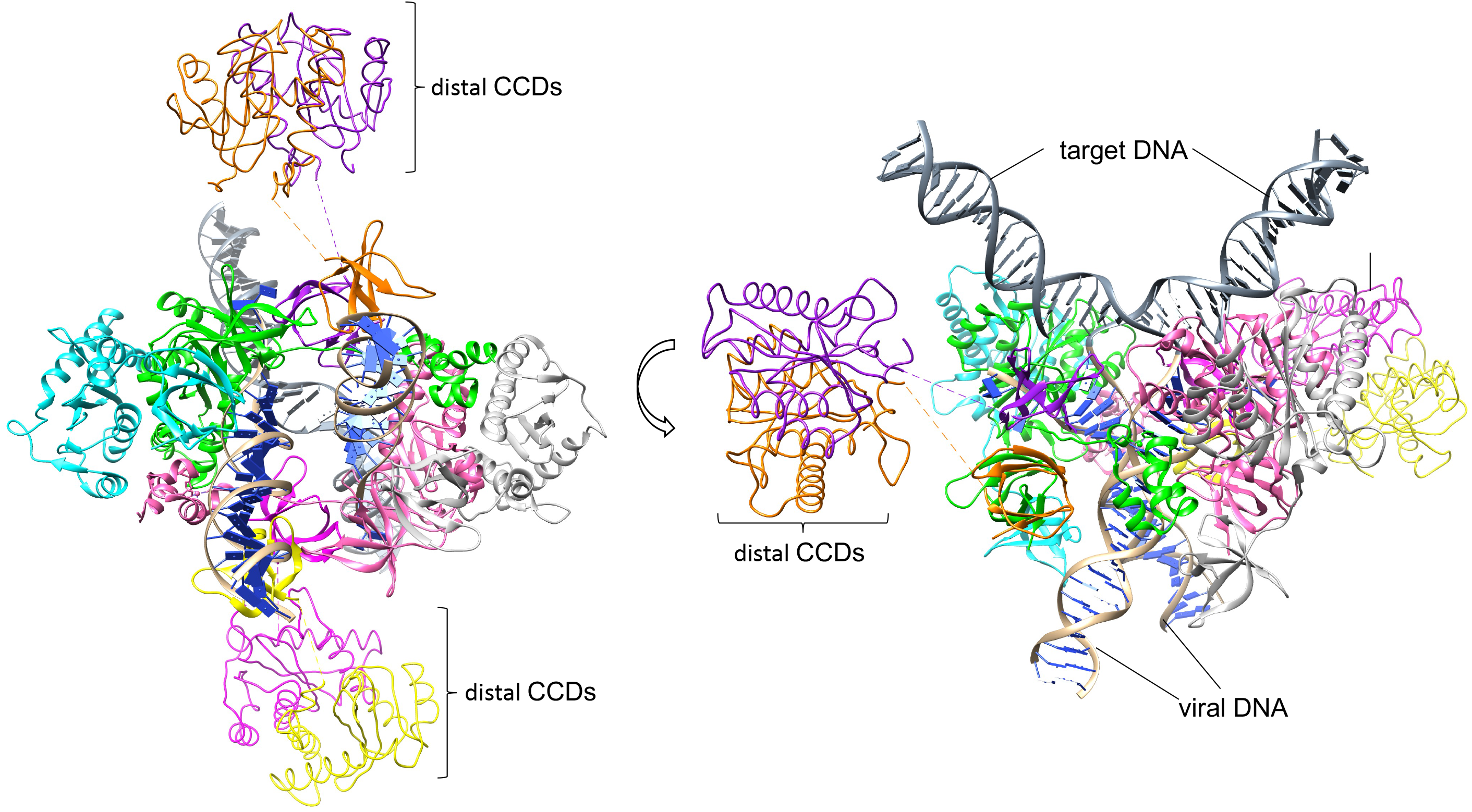
Model of the whole RSV intasome. The backbone of distal CCD subunits were modeled based on their electron densities. The individual subunits follow same coloring scheme as in Fig. 1D.

**Fig S7.**
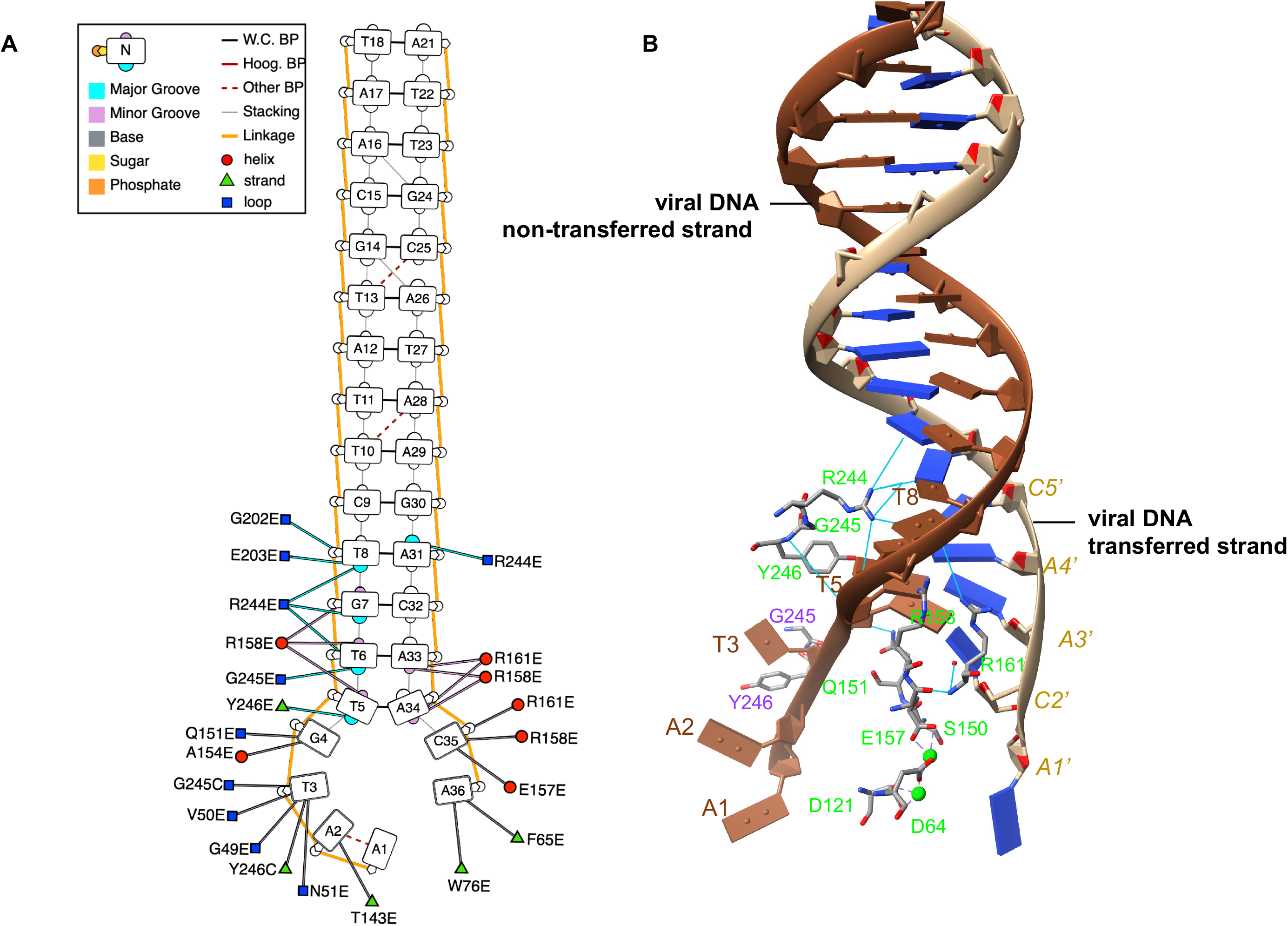
Critical IN-viral DNA contacts in RSV octameric CSC intasome. **A.** Potential nucleotide base-specific interactions including major and minor grove between IN and DNA in the RSV octameric CSC intasome. Analyses were performed using DNAproDB on octameric CSC intasome (PDB 7JN3). Protein residues are labeled with their one letter codes, respective residue number and the protein chain identifier (*e.g.*, R244E indicate R244 of chain E). The protein chains follow the same designation as in Fig. 1D. Schematic shows the potential interactions between IN subunits and nucleotide bases, major and minor grooves. W.C. BP – Watson–Crick bp; Hoog. BP – Hoogsteen bp; Other BP – other bp. **B.** Visualization of select IN-viral DNA interactions in CSC intasome. The select IN residues of proximal and distal subunits interacting with viral DNA are colored by their respective protein chain as in Fig. 1D.

**Fig. S8.**
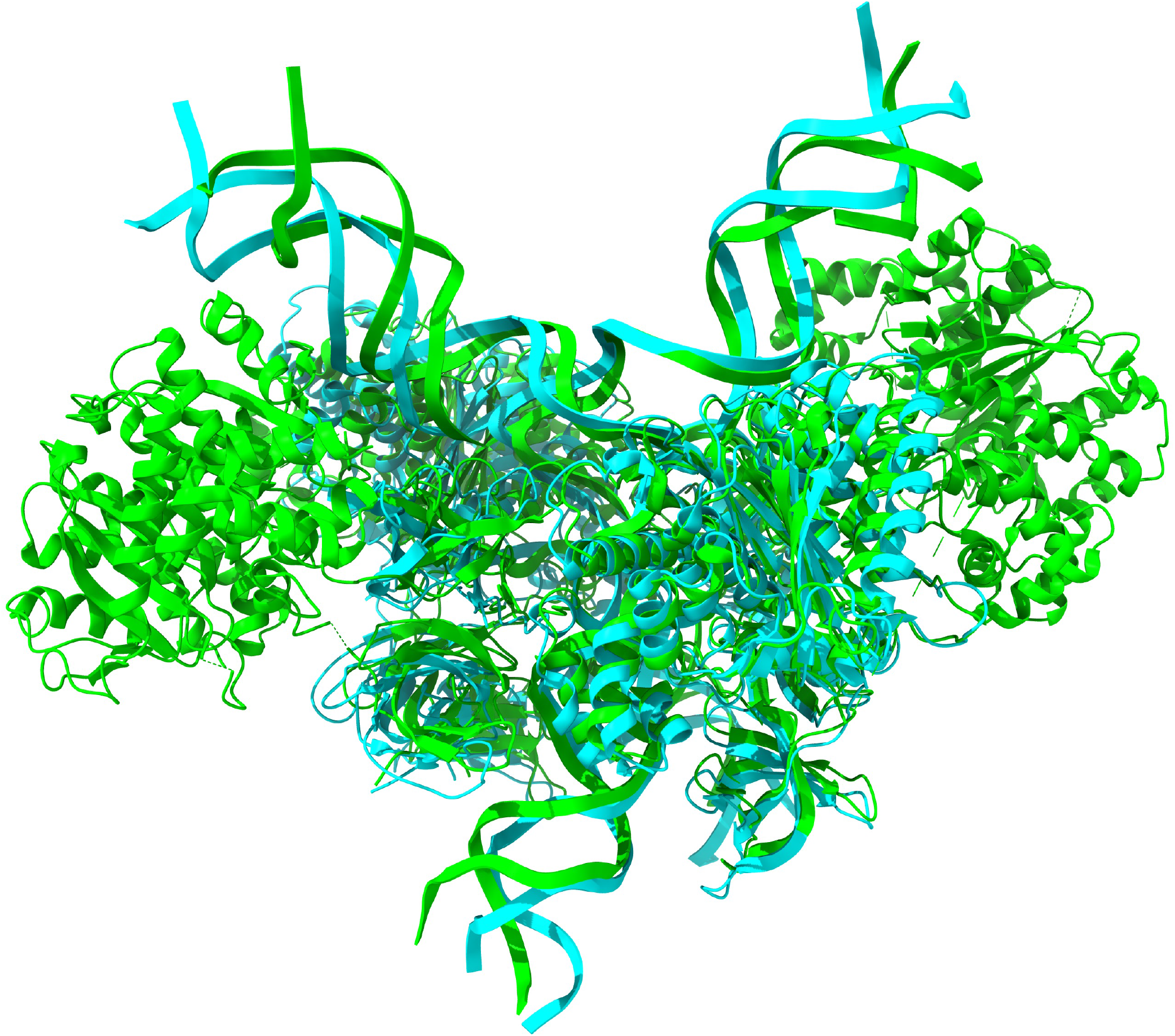
Alignment of the RSV STC structure by cryo-EM (in cyan) and x-ray crystallography (PDB 5EJK) in lime color.

**Supplementary Table 1.**
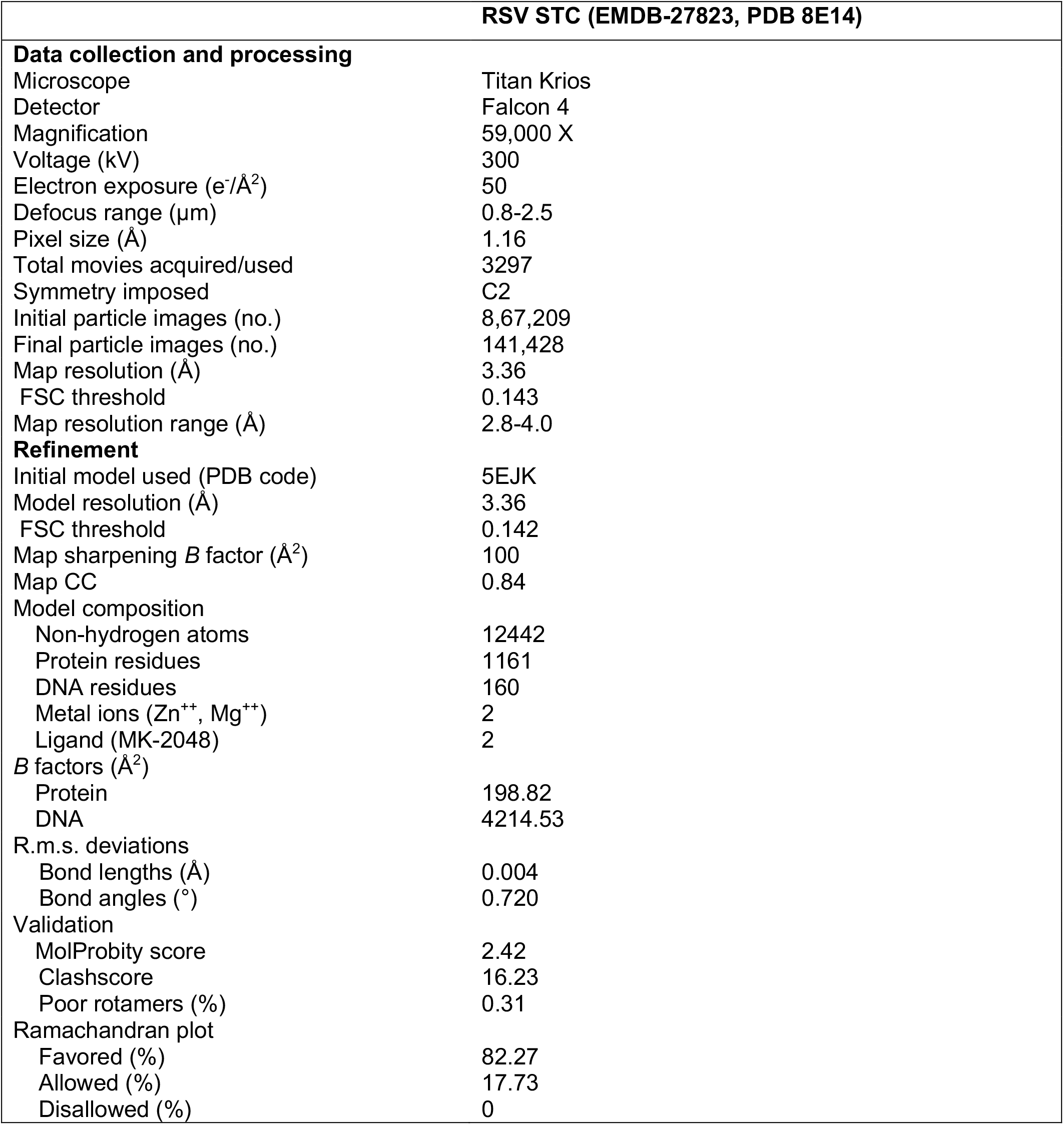
cryo-EM data collection, refinement and validation statistics.

